# Perivascular Space Semi-Automatic Segmentation (PVSSAS): A Tool for Segmenting, Viewing and Editing Perivascular Spaces

**DOI:** 10.1101/2020.11.16.385336

**Authors:** Derek A Smith, Gaurav Verma, Daniel Ranti, Matthew Markowitz, Priti Balchandani, Laurel Morris

## Abstract

**Objective:** In this study, we validate and describe a user-friendly tool for PVS tracing that uses a Frangi-based detection algorithm; which will be made freely available to aid in future clinical and research applications. All PVS detected by the semi-automated method had a match with the manual dataset and 94% of the manual PVS had a match within the semi-automated dataset.

**Methods:** We deployed a Frangi-based filter using a pre-existing Matlab toolbox. The PVSSAS tool pre-processes the images and is optimized for maximum effectiveness in this application. A user-friendly GUI was developed to aid the speed and ease in marking large numbers of PVS across the entire brain at once.

**Results:** Using a tolerance of 0.7 cm, 83% of all PVSs detected by the semi-automated method had a match with the manual dataset and 94% of the manual PVS had a match within the semi-automated dataset. As shown in figure 3, there was generally excellent agreement between the manual and semi-automated markings in any given slice.

**Significance:** The primary benefit of PVSSAS will be time saved marking PVS. Clinical MRI use is likely to become more widespread in the diagnosis, treatment, and study of MS and other degenerative neurological conditions in the coming years. Tools like the one presented here will be invaluable in ensuring that the tracing and quantitative analysis of these PVS does not act as a bottle neck to treatment and further research.

## Introduction

MRI has become a valuable tool in the last several years to aid in the characterization and treatment of neuropsychiatric and neurological diseases, with the advent of ultra-high field imaging (7 T) being of particular note (Wisse et al. 2015; Sladky et al. 2013; Kerchner et al. 2010). 7 T MRI provides exceptionally detailed high resolution structural images, which has enabled *in vivo* study of the progression of neurodegenerative conditions (Liem et al. 2012; Tallantyre et al. 2010; Boutet et al. 2014). MRI has particularly become crucial in describing the presence and progression of perivascular spaces (PVS) (Patankar et al. 2005; Conforti et al. 2014; Groeschel et al. 2006; Kilsdonk et al. 2015; Wuerfel et al. 2008). PVS are small fluid-filled structures that appear on MRI scans in the vicinity of blood vessels. The presence and size of these PVS has been linked to gray and white matter pathology generally, and is of particular interest in a broad range of disorders like multiple sclerosis, cerebral small vessel disease, depression, and dementia (Rouhl et al. 2008; Favaretto et al. 2017; Zhu, Dufouil, et al. 2010; Zhu, Tzourio, et al. 2010; Heier et al. 1989; Patankar et al. 2007; Potter et al. 2015; Thomas et al. 2002). These PVS are believed to be strongly associated with the development and advancement of the disease and describing them *in vivo* is necessary to both clinical treatment of MS, and future research into potential treatments.

Manual detection and tracing of PVS is the current gold standard despite the fact that method is time consuming, prone to rater error/fatigue, and tends to underestimate the number of spaces present in the image. New methods using automated or user-assisted detection algorithms, particularly those using Frangi filters, have been explored and shown to be effective at reducing tracing time while improving accuracy and improving the number of PVS detected in both 3 T and 7 T environments (Dubost et al. 2018; Boespflug et al. 2018; Ballerini et al. 2018; Park et al. 2016; Zhang et al. 2017; Zhang et al. 2016; Hou et al. 2017; Zong et al. 2016). In this study, we validate and describe a user-friendly tool for PVS tracing that uses a Frangi-based detection algorithm with added region growing features; which will be made freely available to aid in future clinical and research applications. 83% of PVS detected by the semi-automated method had a match with the manual dataset and 94% of the manual PVS had a match within the semi-automated dataset.

## Methods

PVSSAS (Perivascular Space Semi-Automatic Segmentation) is a semi-automated tool for segmenting, viewing and editing PVS in the brain.^1^ PVSSAS runs on Matlab and is created using Matlabs inbuilt GUI editor, GUIDE. By default, PVS are segmented using a 2D frangi filter, but any custom segmentation can be used. Frangi filtering is a quick and powerful segmentation technique to segment vessel-shaped objects in 2 and 3 dimensions. Several papers have already applied frangi to 7T PVS segmentation in conjunction with other methods with fair success, although there is a lot of room for improvement (Ballerini et al. 2016; Zong et al. 2016; Hou et al. 2017).

An external frangi filter package made for Matlab is used (https://www.mathworks.com/matlabcentral/fileexchange/24409-hessian-based-frangi-vesselness-filter). Prior to application of the Frangi filter, the PVSSAS program pre-processes the T2-weighted structural images by applying a Gaussian burring function and normalizing the voxel intensity. PVSSAS also pre-processes the white matter mask by filling in small holes, under 200 voxels in size and performing a morphological erosion to remove potential false positives along the gray-white matter boundary. The Frangi filter is applied to the masked white matter region, using an adjustable setting sensitivity parameter (set to 0.5 standard deviations for the presented data). PVSs were also filtered by size to exclude candidates too small to be reliably identified or large but normal anatomical regions of CSF like ventricles. A range of 4 to 300 voxels was selected in consultation with radiologists and researchers on the T2-weighted imaging dataset.

**Figure 1.**
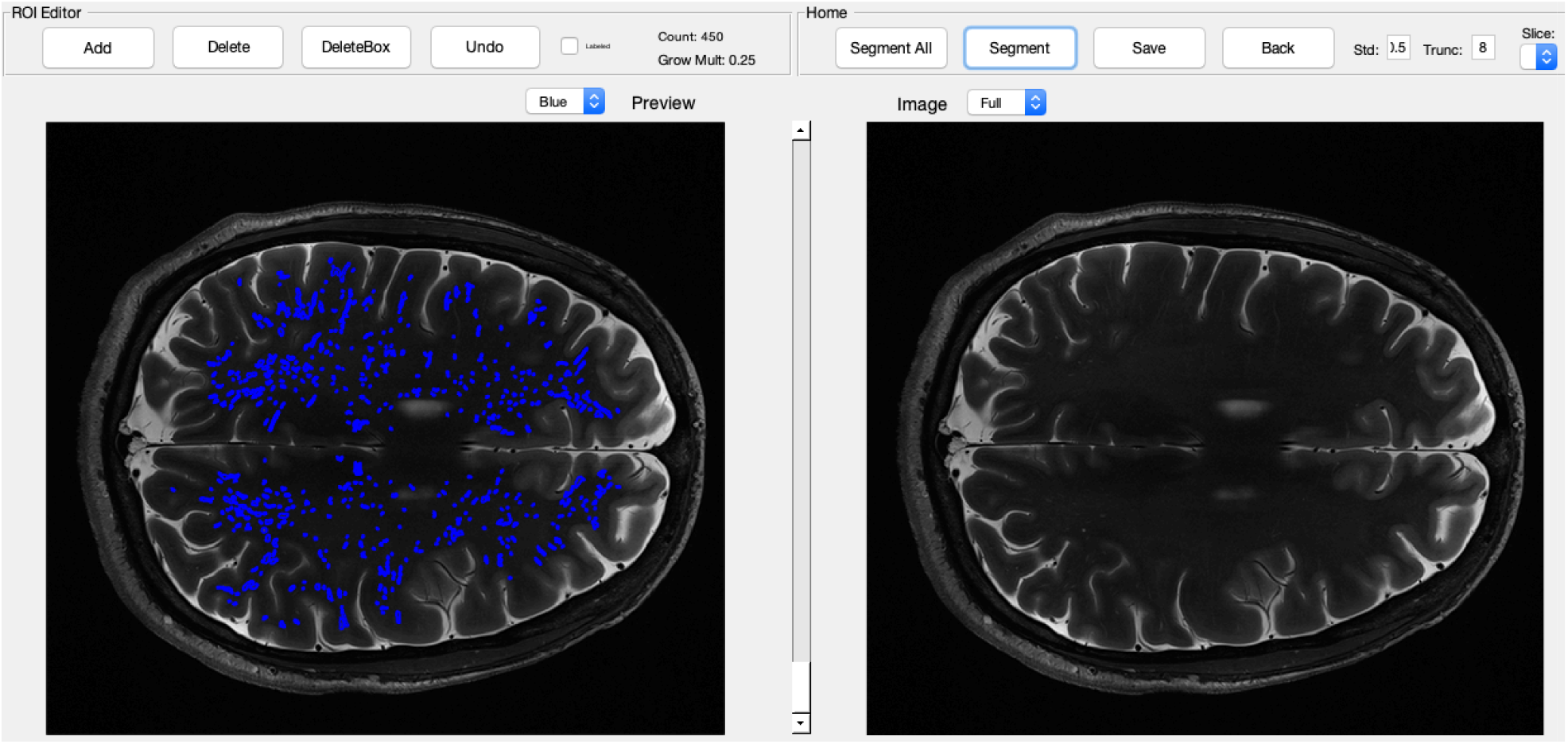
The primary interface for the PVSSAS tool. In the right view panel, the GUI displays raw images for the selected slice. On the toolbar, options are available for segmenting the whole brain, the selected slice, saving tracing masks, or for altering the parameters for the segmentation algorithm. In the left view panel, the completed segmentation can be viewed and edited – a trained reader can add or remove tracings.

Frangi parameters in pvssas_v1-4 were chosen among those that produced fair agreement with several manual markings, however it was noted that these marking under-estimated the number of PVS, and hence newer parameters may be needed. For pvssas_v5, parameters were changed slightly to resemble optimal parameters as determined by Ballerini, L. et al (Ballerini et al. 2018).

### Validation

PVSSAS was validated on a single multiple sclerosis patient, a *male/female* who was n years old at the time of her scan. Our validation of PVSSAS was focused primarily on evaluating the relative speed, accuracy and sensitivity both manual and tool-assisted methods. In the manual method, markings were performed in OsiriX by a trained MD with over 20 hours of experience. PVS were marked in the entire brain, which took approximately 6 hours to manually mark all 54 slices of the scan.^2^ Another rater was then trained to identify PVS on 7 T T2 TSE scans and also trained in use of the PVSSAS tool for 1-2 hours. The rater was told to focus on removing erroneous markings, not adding additional ones. It took approximately 30 minutes for the rater to completely mark the scan using PVSSAS. Separately, we assessed inter-rater reliability with two raters using PVSSAS, who focused on removing erroneous markings, not adding additional ones.

We focused our quantitative comparison of both methods on the sensitivity and specificity of each. This was done due to the fact that the manual tracings performed in OsiriX marked each PVS with a cross-sectional point, rather than as volumes. Therefore, it was not possible to reliably compare other useful measures such as the relative length, average volume or total volumes of each. Sensitivity measures were determined by comparing the total number of PVS found using each method. Certain constraints were enforced: (1) manually traced PVS smaller than a single voxel were ignored, as these would be too small to be reliably detected by the PVSSAS; (2) our method was unable combine PVS that move between slices, so these spaces were counted as separate PVS in both the semi-automated and manually traced methods; (3) all PVS above voxels in size were ignored and this mostly affected semi-automated markings which were unlikely to be PVS; (4) all PVS that fell outside of the white matter mask were not used for comparison, as the semi-automated method is incapable of identifying them. Specificity measures were determined by performing a slice-by-slice comparison between the semi-automated and manual tracings. For each slice the centroid (center voxel) of every tracing was determined using the RegionProps3 command in Matlab, and all centroids within 0.7 cm were considered successful overlap between the manual and semi-automated tracings. This number was determined based on an analysis of the number of PVS overlapping relative to the distance to the nearest match. Above ∼0.7mm, the distance to the closest match rapidly increases (fig. 2).

**Figure 2.**
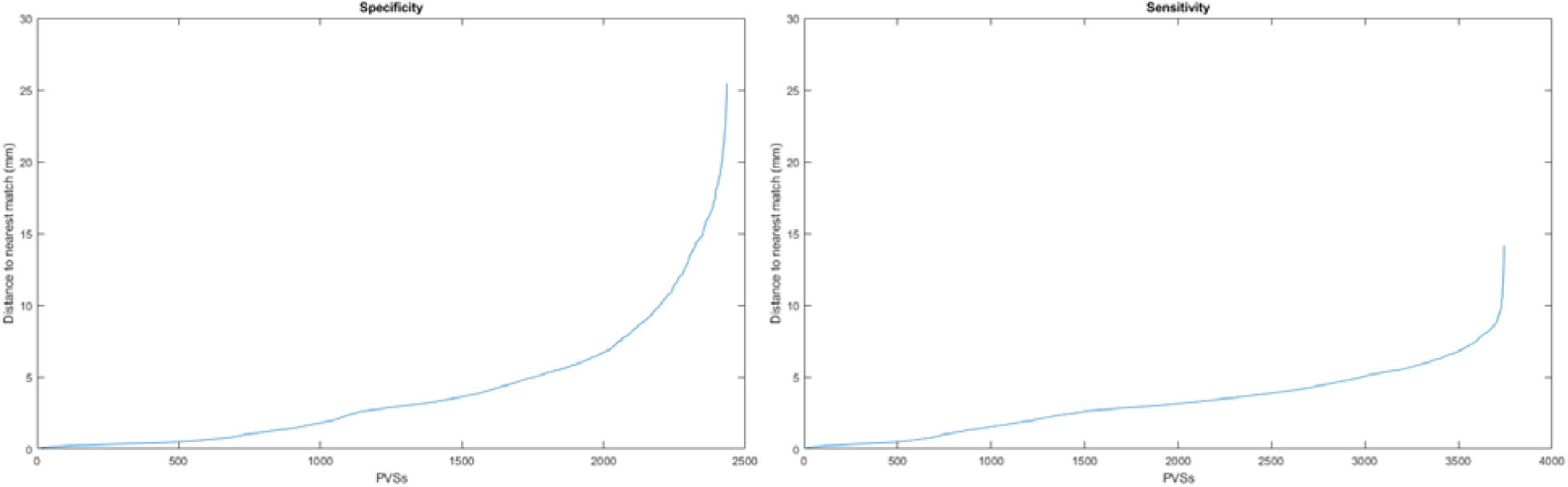
The Specificity (left) plot displays the relationship between the number of PVS found to be overlapping between the manual and semi-automated methods, relative to the distance to the nearest match. The Sensitivity (right) plot shows the number of PVS detected by the semi-automated method in relation to their nearest equivalent neighbors in the manual markings.

**Figure 3&4.**
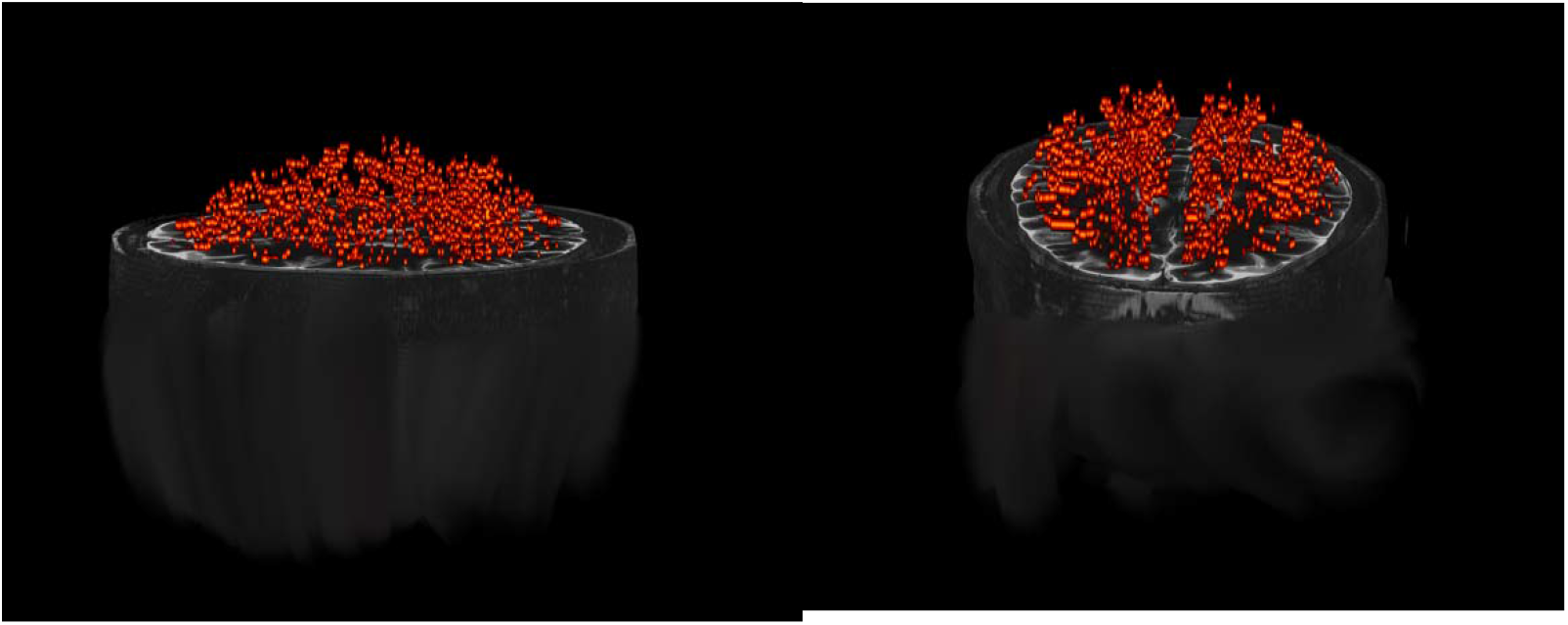
Rendered 3-D view of total PVS found across all slices for one patient.

## Results

The semi-automated method identified a total of 2435 PVS. The manual reader identified 6162 PVS, but these included PVS outside the white matter mask. When applying the white matter mask to the manual segmentation, the PVS count dropped from 6162 to 3743. Using a tolerance of 0.7 cm, 83% of all PVSs detected by the semi-automated method had a match with the manual dataset and 94% of the manual PVS had a match within the semi-automated dataset. As shown in figure 3, there was generally excellent agreement between the manual and semi-automated markings in any given slice.

The inter-rater reliability between the two readers using the semi-automated method was as follows: of the PVS identified by reader 1, 97.8% of the PVS were in the identical location as reader 2, and 99.6% were within 0.8 cm. Of the PVSs identified by reader 2, 99.2% were in the same location as reader 1, and 99.9% were within 0.8 cm. Reader 1 identified 1680 PVSs, and reader 2 identified 1656 PVS. The Dice score between the two readers segmentations was 0.9914.

## Discussion

The primary benefit of PVSSAS will be time saved marking PVS. Using a tool such as PVSSAS, one rater with only a moderate amount of additional training to become familiarized with the UI can perform the same amount of total brain markings that currently requires multiple raters, at the combined total of 5-10+ man-hours per rater. Clinical MRI use is likely to become more widespread in the diagnosis, treatment, and study of MS and other degenerative neurological conditions in the coming years. Tools like the one presented here will be invaluable in ensuring that the tracing and quantitative analysis of these PVS does not act as a bottle neck to treatment and further research.

PVSSAS also provides valuable information that is currently not provided from other manual tracing programs, such as OsiriX. Typically, the process for marking an entire brain worth of PVS is very time consuming, and it is often unreasonable to expect raters to individually trace the entire volume of each PVS in an entire brain. However, the method used by PVSSAS automatically captures volume information for PVS, which could allow for more varied analysis of these spaces.

### Limitations

In the current implementation of the PVSSAS there are certain restraints on both the sensitivity and specificity. The frangi filter used as the basis for PVSSAS is designed to look for objects at least 4 voxels in size. Similarly, PVSSAS has a maximum size of 100 voxels, although occasionally multiple traced objects in inadvertently overlap. Often this is easily spotted by raters, although it can affect the ability to reliably calculate the volume and centroid of certain PVS in a set of tracings. We recommend that users discount all tracings below 4 voxels and above 100 voxels as being erroneously marked, and focus on these areas as a good starting point for manually marking the remainder of spaces. Even in spite of these limitation, the relatively high accuracy and sensitivity of the tool means that it provides a reliable starting point for a rater to finish a tracing much faster than a purely manual method.

## Acknowledgements

Thank you to Ameen Al Qadi for his work in providing 3-D renderings for Figure’s 3 and 4.

## Footnotes

1 PVSSAS is available for download at: https://github.com/smithd37/pvssas

2 Note: this includes certain areas that fall outside of the white-matter mask, and are therefore undetectable to the PVSSAS tool.

